# Triggered Temptations: A New Procedure to Compare Reward-Seeking Behaviour Induced by Discriminative and Conditioned Stimuli in Rats

**DOI:** 10.1101/2024.12.06.627237

**Authors:** Mandy Rita LeCocq, Shaghayegh Najafipashaki, Domiziana Casale, Isabel Laplante, Anne-Noël Samaha

**Affiliations:** Department of Pharmacology and Physiology, Faculty of Medicine, Université de Montréal, Montréal, QC, Canada; Department of Psychology, Faculty of Medicine, Université de Montréal, Montréal, QC, Canada; Neural Signaling and Circuitry Research Group, Faculty of Medicine, Université de Montréal; Centre for Interdisciplinary Research on the Brain and Learning (CIRCA), Université de Montréal, Montréal, QC, Canada; Centre for Biomedical Innovation (CIB), Université de Montréal, Montréal, QC, Canada; Centre for Studies in Behavioural Neurobiology, Montreal, QC, Canada

**Keywords:** Sucrose self-administration, Appetitive, Reward, Pavlovian conditioning, Instrumental conditioning, Conditioned reinforcement

## Abstract

**Rationale:** Environmental cues guide animals towards resources vital for survival but can also drive maladaptive reward-seeking behaviours, as in gambling and eating disorders. While conditioned stimuli (CSs) are paired with reward delivery after reward-seeking actions, discriminative stimuli (DSs) signal reward availability independent of behaviour.

**Objective:** We introduce a procedure to compare CS and DS effects on reward-seeking behaviour, in the same subjects within a single session.

**Methods:** Female and male Sprague-Dawley rats learned to self-administer sucrose. During each session, DS+ trials signaled that lever pressing would produce sucrose paired with a CS+, and DS−trials signaled no sucrose and a CS−. Next, in the absence of sucrose, we assessed the ability of the cues to *i)* reinforce lever pressing and *ii)* increase sucrose seeking when presented response-independently. We also assessed the effects of the mGlu_2/3_ receptor agonist LY379268 and d-amphetamine on cue-induced sucrose seeking.

**Results:** By the end of self-administration training, lever pressing peaked during DS+ trials and dropped during DS−trials. The DS+ was a conditioned reinforcer of sucrose seeking in both sexes, whereas the CS+ was more effective in males. Response-independent presentations of the DS+ invigorated sucrose seeking in both sexes, whereas the CS+ was effective only in males. LY379268 suppressed DS+-triggered sucrose seeking in females, with no effect in males. D-amphetamine enhanced sucrose seeking non-specifically across cue conditions in males, with no effect in females.

**Conclusions:** Our new trial-based procedure can be used to identify unique and similar mechanisms underlying DS and CS influences on appetitive behaviour.

## Introduction

Animals rely on environmental cues to guide their actions toward rewards necessary for survival and fitness, including food and water (Balleine & Dickinson, 1998). Through associative learning, such cues gain motivational salience and the power to influence appetitive behaviour (Robinson & Berridge, 1993). In parallel, the same mechanisms that underlie adaptive cue-guided reward seeking can also promote maladaptive reward seeking that contributes to gambling (Barrus et al., 2016; Kushner et al., 2008), eating (Boswell & Kober, 2016) and addiction (Obrien et al., 1992; Shulman, 1989; Vafaie & Kober, 2022) disorders.

Reward cues include conditioned stimuli (CSs) and discriminative stimuli (DSs). In instrumental settings, conditioned stimuli (CSs) are paired with reward delivery and their presentation is dependent on reward-seeking actions (Bindra, 1978; Holland & Rescorla, 1975; Weiss, 2005). Therefore, CSs and the neural substrates they recruit operate after reward seeking is initiated. For example, the taste of fruit is a CS that can maintain and invigorate ongoing eating. In comparison, discriminative stimuli (DSs) signal reward availability or unavailability, and their presentation is typically independent of any reward-seeking action (Colwill & Rescorla, 1988). For instance, the sight of a patch of bright red berries can prompt a bird to redirect its flight path to obtain the ripe food. In contrast, the sight of barren land can indicate the unavailability of food, resulting in the bird staying on its course. DSs are unique as they can occur before, during, and after reward seeking, such that DSs can both trigger and maintain reward seeking (Falk, 1994; Falk & Lau, 1995; McFarland & Ettenberg, 1997).

CSs and DSs influence reward-seeking behaviour in similar and distinct ways. When presented contingent upon responding, both DSs and CSs maintain reward-seeking actions in the absence of the primary reward (Mackintosh, 1974; Milton & Everitt, 2010; Ndiaye et al., 2024; Nie & Janak, 2003; Robinson & Berridge, 1993; Servonnet et al., 2020; Shaham et al., 2003; Stewart, 1984). However, when presented response-independently, CSs are relatively ineffective at increasing reward-seeking behaviour, while DSs are particularly effective even after extended abstinence periods or repeated testing (Ciccocioppo et al., 2001; Deroche-Gamonet et al., 2002; Di Ciano & Everitt, 2003; Madangopal et al., 2019, 2021; Ndiaye et al., 2024; Pitchers et al., 2015; Weissenborn et al., 1995; Yun & Fields, 2003).

Previous studies have informed on how CSs and DSs influence reward-seeking behaviour in animal models; however, there are two main limitations to this work. First, DSs signalling reward availability (DS+) and unavailability (DS−) are often presented as contextual stimuli in separate training sessions (Ciccocioppo et al., 2004; Laque et al., 2019; Martin-Fardon & Weiss, 2017; Suto et al., 2016; Weiss et al., 2000; Zhao et al., 2006). Although contexts can be powerful DSs (Fuchs et al., 2009; Fuchs, Evans, Ledford, et al., 2005; Xie et al., 2009), DSs can also be experienced as discrete events, such as when a person with a drug addiction disorder runs into a drug-using partner (DS+) by happenstance. Therefore, the effects of discrete, trial-based DS presentations are also important to model in preclinical studies (see also Madangopal et al., 2019, 2021). Second, the effects of CSs and DSs on reward seeking are often compared in tests presenting the DS and CS in combination (Ciccocioppo et al., 2001; Collins et al., 2023) or in different groups of animals across different experimental conditions (Collins et al., 2023; de Wit & Stewart, 1981; Di Ciano & Everitt, 2003; Grimm et al., 2002; Lu et al., 2004; Milton & Everitt, 2010; Pitchers et al., 2017; Shalev et al., 2002; Weiss, 2005; Yun & Fields, 2003). This makes it challenging to isolate the unique influences of DSs and CSs on reward seeking behaviour in the same individuals.

Here we developed a novel discrimination procedure with two key advantages. First, during discrimination training, the DS+ and the DS−are presented within the same session and environmental context, in trial-based fashion. Second, our new procedure allows one to distinguish and directly compare the respective ability of CSs and DSs to *i)* act as conditioned reinforcers of reward-seeking actions and *ii)* spur increases in reward-seeking actions, all in the same subjects with each process evaluated in a single session. Finally, considering that the selective metabotropic glutamate type II receptor agonist suppresses the response to reward cues (Bäckström & Hyytiä, 2005; Baptista & Weiss, 2004), and that D-amphetamine enhances this response (Parkinson et al., 1999; Taylor & Robbins, 1984; Wyvell & Berridge, 2000), we also examined the effects of LY379268 and D-amphetamine on reward seeking mediated by a DS and a CS.

## Methodology

### Subjects

Female (*n* = 15; 11 weeks on arrival) and male (*n* = 15; 8 weeks on arrival) Sprague-Dawley rats (Charles River Laboratories, Raleigh, North Carolina, Barrier R04) were individually housed in a climate-controlled animal facility with ad libitum access to water and food unless specified otherwise. The facility was maintained at 21°C, under a 12-hour reverse light/dark cycle with lights OFF at 8:30 AM. All behavioural procedures were conducted during the dark phase. Housing cages included Betachip bedding and a chew toy (Nylabones, Bio-Serv, New Jersey U.S.A). After a 72-hour acclimation period, the experimenters handled the rats daily for five days. All experimental protocols adhered to the guidelines of the Canadian Council on Animal Care and were approved by the Université de Montréal Animal Research Ethics Committee.

The present rats were first used in a separate experiment involving optogenetic stimulation of ChR2-expressing neurons in the lateral orbitofrontal cortex. Thus, under isoflurane anesthesia (5% for induction, 1%–2% for maintenance), the rats received both bilateral adeno-associated virus infusions [AAV2/7-CamKIIa-hChR2(H134R)-eYFP or AAV2/7-CamKII-eYFP] and optic fiber implants into the lateral orbitofrontal cortex. This surgery occurred before the behavioural training described below. The present experiment only includes data from eYFP- and ChR2-expressing rats during sessions where they did not receive laser delivery.

### Conditioning chambers

Behavioural training and testing were conducted in 8 operant chambers (Med Associates, St. Albans, VT), placed within light- and sound-attenuating boxes equipped with ventilation fans. Each chamber included a clicker, a tone generator (75 dB, 2900 Hz), a liquid dispenser connected to a 20-ml syringe containing 10% *w*/*v* sucrose solution, and two retractable levers. During self-administration sessions, pressing the inactive lever (counterbalanced across left- and right-side levers) had no programmed consequence, and pressing the active lever produced 0.1 ml sucrose solution delivered into a magazine equipped with two infrared photobeams to record magazine entries. Each chamber had 4 cue lights; one above each lever, two on the wall opposite to the levers positioned respectively, at the bottom left corner and top center. Finally, the chambers had two infrared photobeams, evenly spaced 3.5 cm above the grid floor to quantify horizontal locomotor activity. All data were recorded and analyzed using a PC running Med-PC IV software.

### Drugs

LY379268 (Tocris, CAT#2453; 0.3 and 1 mg/kg) was dissolved in 0.9% saline. The solution was heated to 45 ºC and then sonicated for 10 minutes. The pH was then adjusted to 7 with 10 N NaOH. D-amphetamine (Sigma-Aldrich, Dorset, UK; CAT# A5880) was dissolved in 0.9% saline. Both drugs were administered subcutaneously (s.c.) in a volume of 1 ml/kg. A within-subjects design was used such that all rats got saline vehicle, 0.3 and 1 mg/kg LY379268, and 0.75 mg/kg D-amphetamine.

### Sucrose habituation in the home cage

To facilitate subsequent instrumental sucrose self-administration, rats were first habituated to liquid sucrose in their home cage. For 48 h, rats had access to a bottle containing a 10% *w*/*v* sucrose solution, available alongside their regular water bottle.

### Water restriction

Following sucrose habituation, access to water was restricted to promote subsequent sucrose self-administration. For the first 4 days, rats had access to water for 6 h/day, then 4 h/day for 3 days, and finally, 2 h/day until the end of the study. Rats were given access to water for at least 1 h after daily behavioural training or testing sessions.

### Magazine training

To familiarize rats with retrieving sucrose from the magazine, they received 2 magazine training sessions (30 min/session). During each session, 30 sucrose reinforcers (0.1 ml each) were delivered on a variable interval (VI) 45-s schedule of reinforcement. Magazines were checked at the end of each session to verify that rats retrieved the sucrose.

### Autoshaping

When a conditioned stimulus (CS) predicts delivery of an unconditioned stimulus (US), rats can develop different Pavlovian conditioned approach phenotypes: a sign-tracking (approach to the CS), goal-tracking (approach to the site of reward delivery), or intermediate (approach to both the CS and site of reward delivery) phenotype. Rats with a sign-tracking phenotype have been shown to respond more to reward-associated CSs whereas rats with goal-tracking phenotype respond more to reward-associated DSs (Flagel et al., 2011; Pitchers et al., 2017; Saunders et al., 2014).

To account for these potential individual differences, after magazine training, rats received 6 Pavlovian autoshaping sessions (30 min/session; 1 session per day) to determine their Pavlovian conditioned approach phenotype. Fig. 1A illustrates the Pavlovian autoshaping procedure. During autoshaping sessions, a lever CS extended for 8 s and was followed by delivery of 0.1 ml of sucrose solution (the US), on a VI 30-s schedule with a total of 40 reinforcers/session. This lever was the active lever used in following experimental phases. A response bias score was calculated to determine conditioned approach phenotype, using the following equation: 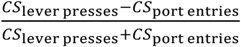 (Flagel et al., 2007; Meyer et al., 2012; Robinson & Flagel, 2009). Based on the average score of the two last sessions, rats were classified as sign-trackers (response bias between +0.5 and +1), goal-trackers (response bias between −0.5 and −1) or intermediates (response bias between −0.49 and +0.49).

**Figure 1.**
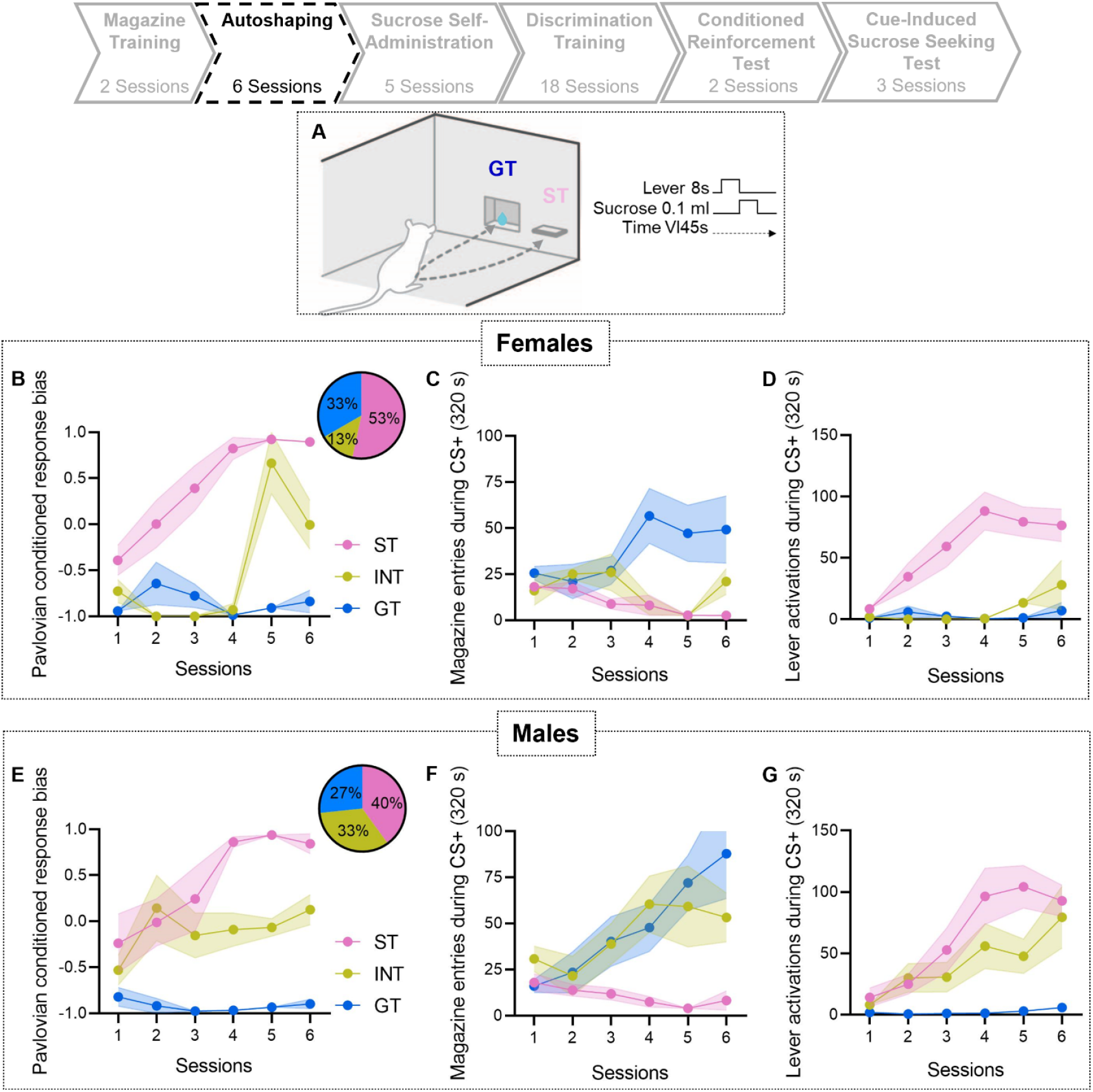
Acquisition of Pavlovian conditioned responding in female and male rats across autoshaping sessions. Schematic of the Pavlovian conditioned autoshaping procedure. Average (± SEM) Pavlovian conditioned response bias scores across autoshaping sessions for sign-trackers (pink; scores between +0.5 and +1.0), goal-trackers (blue; scores between −0.5 and −1.0), and intermediates (yellow; scores between −0.49 and 0.49) in **(B)** females and **(E)** males. Insets in **(B)** and **(E)** represent percent of rats with each phenotype. Average (± SEM) magazine entries in **(C)** females and **(F)** males, and lever activations in **(D)** females and **(G)** males, during conditioned stimulus (CS+) presentation. Schematic in **(A)** adapted from Servonnet et al. (2023).

### Sucrose self-administration training

Following autoshaping, rats were trained to self-administer sucrose. Daily sessions lasted 1 h or until 100 reinforcers were earned. No cues were presented during these sessions. Sessions began with both levers extending. Rats first learned to press the active lever to earn sucrose on a fixed ratio one (FR1) schedule of reinforcement. Once rats earned 20 reinforcers and pressed twice as much on the active vs. inactive lever for two consecutive sessions, they received sessions on a FR3 schedule. Once rats earned 30 reinforcers while maintaining a 2:1 ratio of active to inactive lever presses for two consecutive sessions, they transitioned to discrimination training. All rats received three FR1 sessions and two FR3 sessions.

### Sucrose self-administration under the control of discriminative stimuli

#### Discrimination training with DSs presented in predictable order

After acquiring the sucrose self-administration task, rats received discrimination training (90 min/daily session). Fig. 2A illustrates the discrimination procedure. During these sessions, discriminative stimuli signaled sucrose availability (DS+) or unavailability (DS−). The DS+ was the cue light positioned at the top center of the back wall. The DS−was the cue light above the active or inactive lever (counterbalanced across rats). The CS+ and CS−were audiovisual stimuli. For half of the rats, the CS+ was a 5-s presentation of the cue light above the active or inactive lever paired with a tone, and the CS−was a 5-s presentation of the cue light at the bottom left corner of the back wall, paired with a clicker sound. For the other half, the CS+ and CS−modalities were reversed.

**Figure 2.**
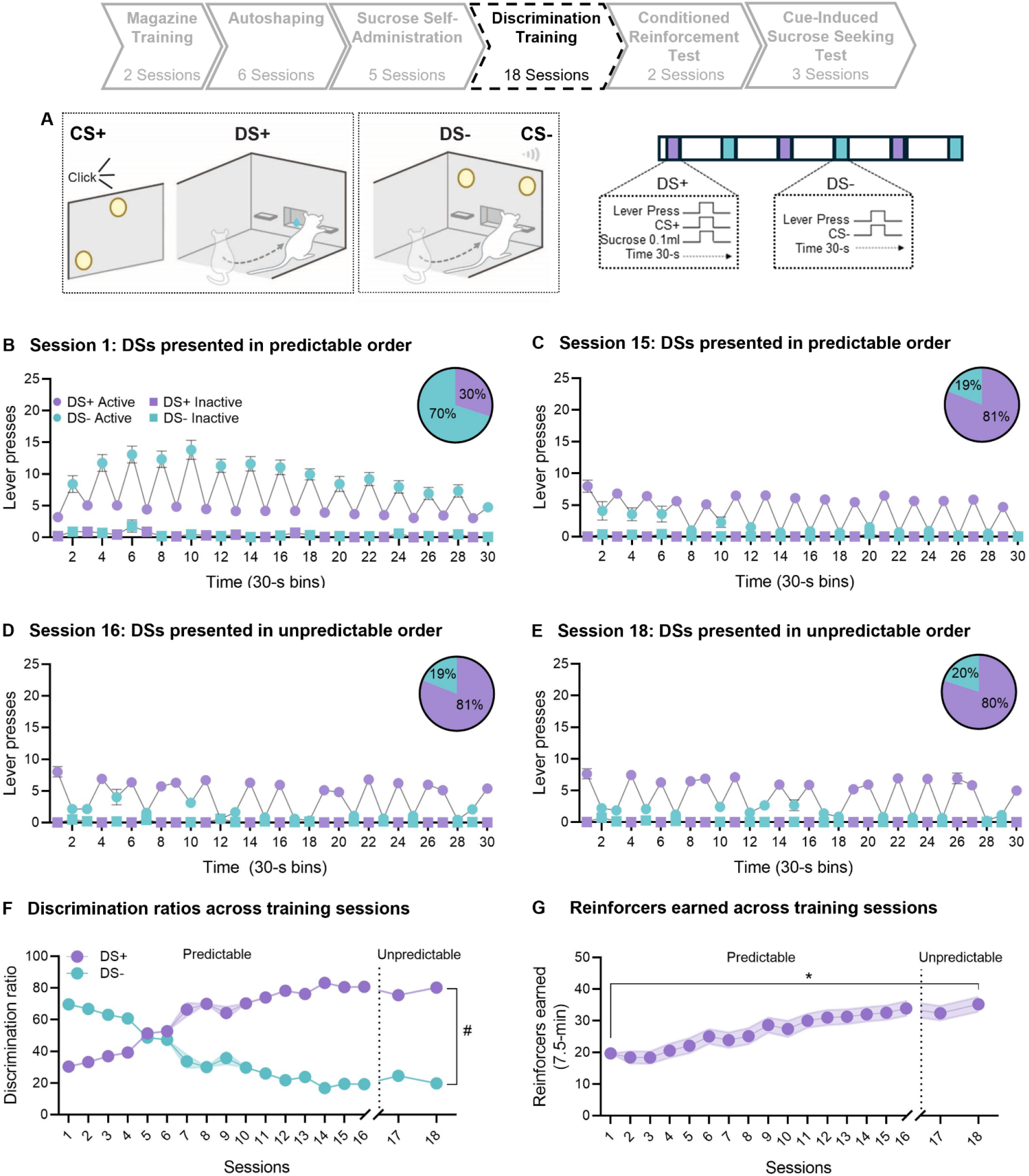
Sucrose self-administration is under the control of discriminative stimuli signaling reinforcer availability (DS+) and unavailability (DS−). **(A)** Schematic of the discrimination procedure. Average (± SEM) active (circle) and inactive (square) lever pressing during DS+ (purple) and DS−(turquoise) presentations during **(B)** session 1 and **(C)** session 15, where the DSs were presented in predictable order, and during **(D)** session 16 and **(E)** session 18, where the DSs were presented in unpredictable order. Insets in **(B-E)** represent percent of active lever presses made during the DS+ and DS−. Average (± SEM) **(F)** discrimination ratios and **(G)** sucrose reinforcers earned across sessions. Data are pooled across female and male rats. Schematic in **(A)** adapted from Servonnet et al. (2023).

During the first 15 discrimination training sessions, the DS+ and DS−were presented in a predictable order, separated by a 2-min inter-trial interval (ITI). Each session began with the levers remaining retracted and the DS+ being presented for 10 s. This was followed by extension of both levers into the cage, and the DS+ staying on for an additional 30 s. Active lever presses during DS+ trials produced sucrose delivery under a random ratio 3 schedule of reinforcement. Each sucrose delivery was paired with a 5-s CS+ presentation. Lever pressing during CS+ presentations had no programmed consequences. The 30-s DS+ trial was followed by the ITI, during which all cues were turned off and the levers retracted. Next, the DS−was presented for 10 s, followed by levers extending and the DS−remaining on for another 30 s. During DS−trials, sucrose was not available and active lever presses produced a 5-s CS−presentation under a random ratio 3 schedule. Lever pressing during CS−presentations had no programmed consequences. This sequence of alternating DS presentations continued until rats received 15 presentations of each DS type.

Discrimination learning was measured using a DS+ discrimination ratio 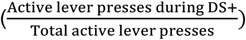 representing the percentage of active lever presses made during the DS+ relative to all active lever presses. We also computed a DS−discrimination ratio 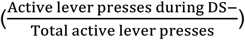. Rats that had a DS+ discrimination ratio of ≥ 60% and earned ≥ 6 reinforcers/session on two consecutive sessions were considered to have learned the discrimination task. By session 8, three males and six females had not met these criteria. These rats were given 2 additional sucrose self-administration sessions (1 h/session) during which the DS+ was presented continuously, and active lever presses resulted in sucrose delivery paired with a 5-s CS+ presentation, under FR3. These rats were then returned to discrimination training. All 30 rats then met the acquisition criteria by session 15.

#### Discrimination training with DSs presented in unpredictable order

During the first 15 sessions, the DS+ and DS−were presented in a predictable order, and the ITI durations were held constant. Under these conditions, the peaks in sucrose self-administration during DS+ trials and the dips in responding during DS−trials could indicate that the DSs controlled sucrose self-administration behaviour, or alternatively, that the rats were keeping time. To address this, rats received 3 additional discrimination training sessions during which the DS+ and DS−were presented in pseudo-random order on a VI 2-min schedule (possible intervals: 60, 90, 120, 150, 180 s), with each DS presented first on 50% of the sessions. If sucrose self-administration is under the control of the DSs, responding should still peak during DS+ periods and dip during DS−periods. Note that each test described next was preceded by two additional discrimination training sessions with DSs presented in unpredictable order.

### Tests for conditioned reinforcement

We then evaluated the conditioned reinforcing properties of the DSs and CSs by measuring the ability of the cues to reinforce lever pressing in the absence of sucrose (as illustrated in Fig. 3A). Conditioned reinforcement tests (50 min/test), began with presentation of the DS+, DS−, CS+, CS− or a No Cue condition for 10 s. After this 10-s cue prime, the cues were turned off and the levers extended for 30 s, during which pressing the active lever produced a 5-s presentation of the same cue condition presented during the 10-s cue prime. Each condition was presented 4 times in pseudo-randomized order, separated by a VI 2-min ITI during which levers were retracted.

**Figure 3:**
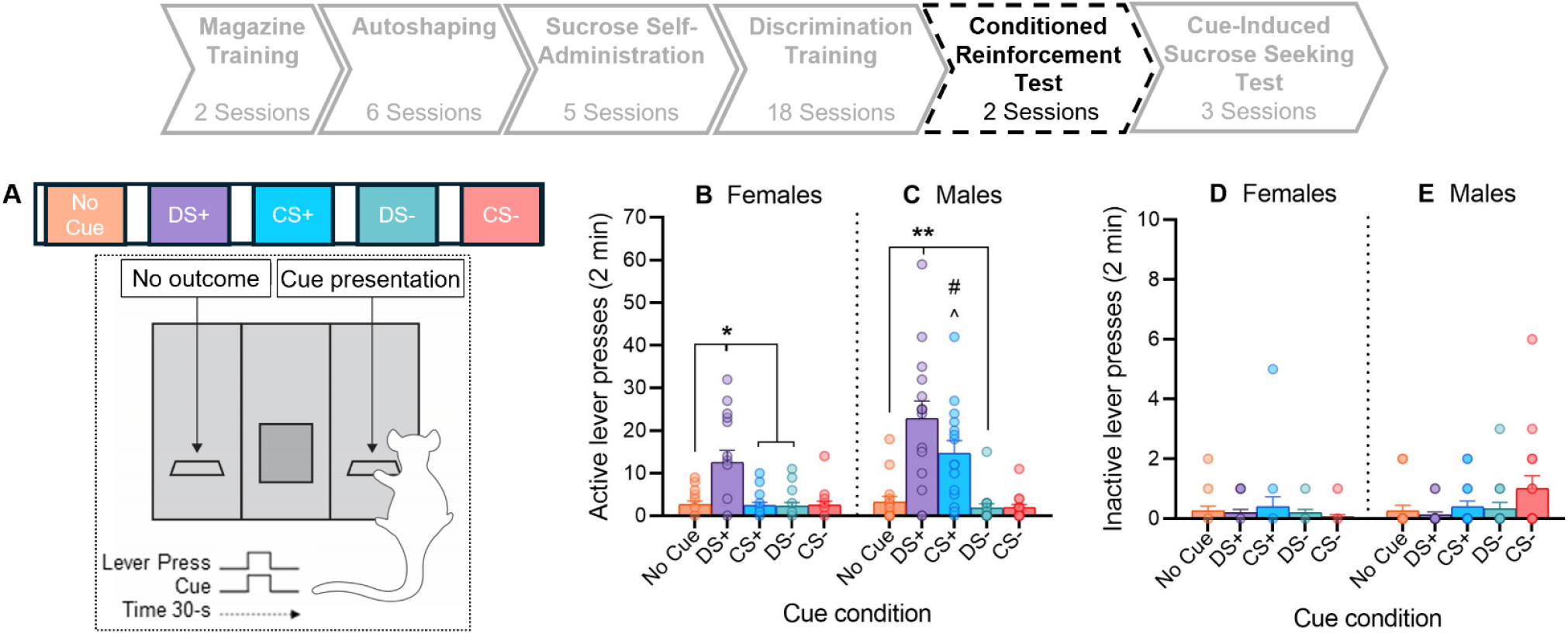
Conditioned reinforcing properties of the DS+ and the CS+ when all cue types were tested in a single session. **(A)** Schematic of the conditioned reinforcement test procedure. Rats could press the active lever to obtain DS+, CS+, DS− or CS−presentations, or no cue. Average (± SEM) active lever presses in **(B)** females and **(C)** males. Average (± SEM) inactive lever presses in **(D)** females and **(E)** males. # CS+ active lever pressing in Males > Females. ^ CS+ > CS−, and No Cue. Schematic in **(A)** adapted from Servonnet et al. (2020).

In the conditioned reinforcement test above, rats could respond for all cue types in the same session. To extend our findings, we gave all rats 2 additional conditioned reinforcement tests during which they could respond for either the DS+ or the CS+ on separate sessions (as illustrated in Fig. 4A). Each test included, respectively, DS+ and No Cue trials, or CS+ and No Cue trials, as described above. Each rat received both test sessions (20 min/session) in counterbalanced order. Two male rats accidentally received the wrong MedPC program and so their data are excluded for these 2 tests.

**Figure 4:**
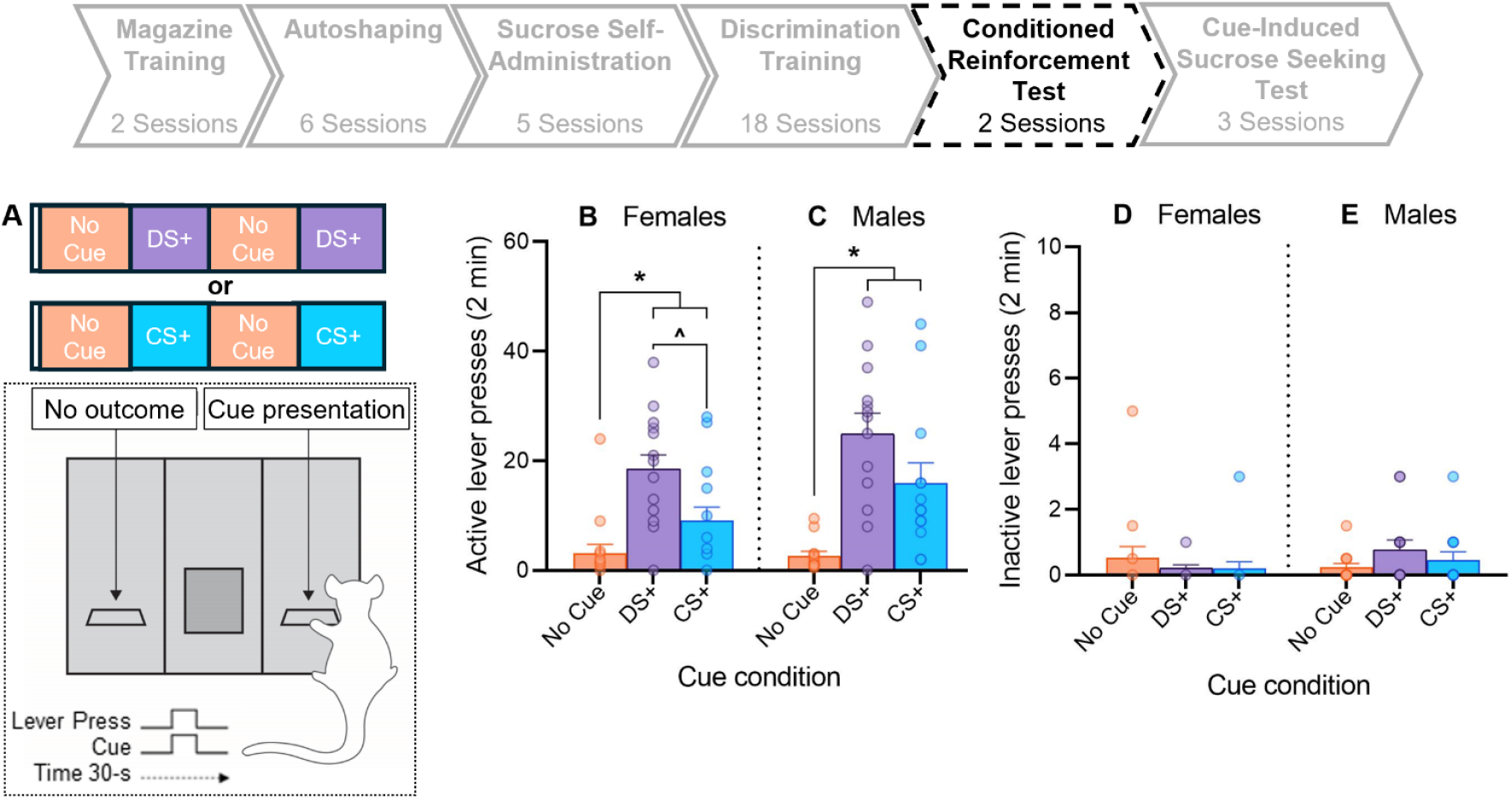
Conditioned reinforcing properties of the DS+ and CS+ when these cues were tested in separate sessions. **(A)** Schematic of the conditioned reinforcement test procedure. On one test session, rats could press the active lever to obtain the DS+ or no cue. On another test session, rats could press the active lever to obtain the CS+ or no cue. Average (± SEM) active lever presses in females and **(C)** males. Average (± SEM) inactive lever presses in **(D)** females and **(E)** males. Schematic in **(A)** adapted from Servonnet et al. (2020).

### Cue-induced sucrose seeking

Following conditioned reinforcement tests, we assessed the ability of the cues to trigger increases in sucrose-seeking behaviour in the absence of sucrose. Cue-induced sucrose seeking tests (50 min; illustrated in Fig. 5A) began with a 5-s presentation of the DS+, DS−, CS+, CS− or No Cue condition. After this 5-s period, the cue (or No Cue condition) stayed on for an additional 30-s, during which the levers were inserted and active or inactive lever pressing had no programmed consequences. Each cue condition was presented 4 times in pseudo-randomized order, separated by a VI 2-min ITI.

**Figure 5:**
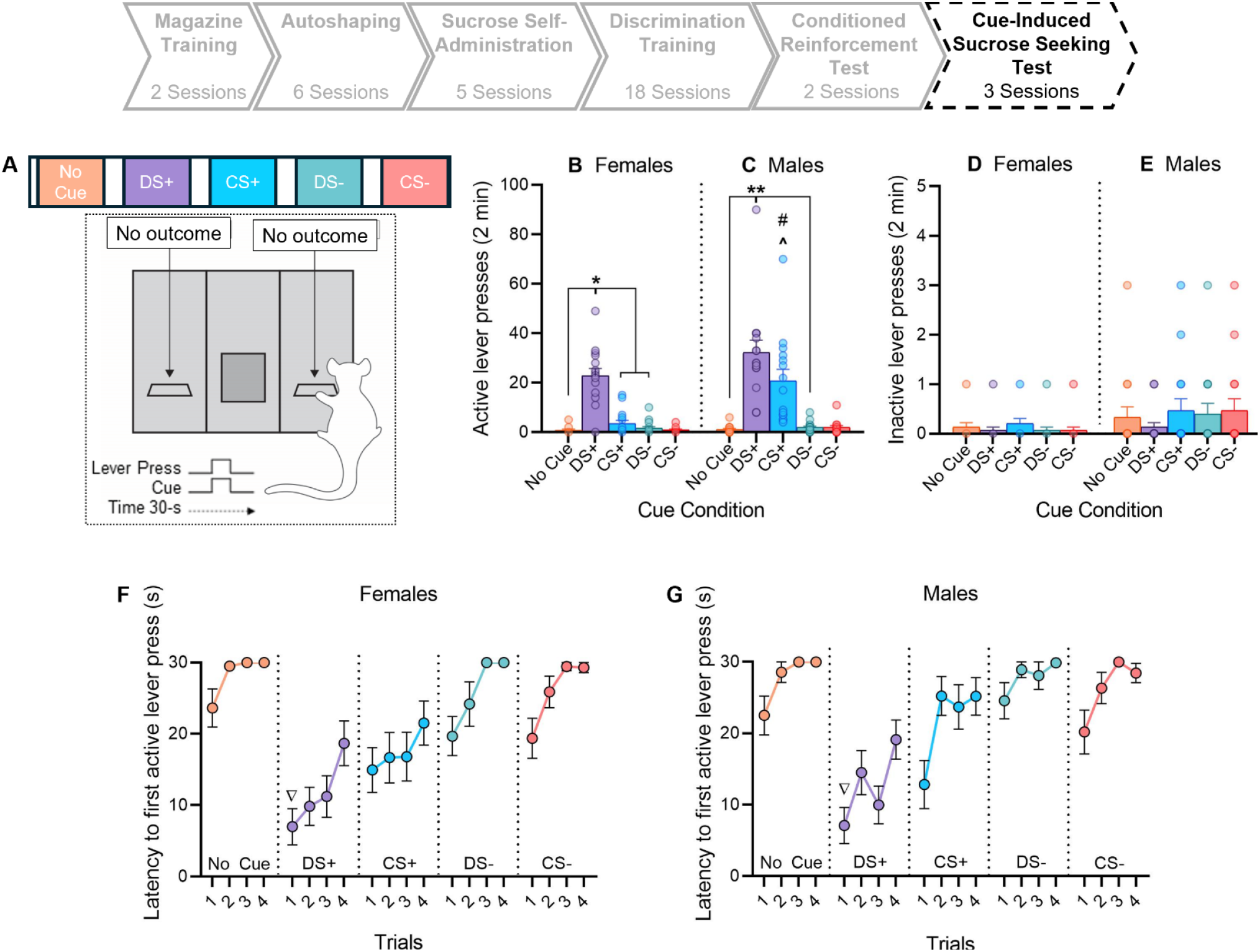
Cue-induced sucrose seeking. **(A)** Schematic of the cue-induced sucrose seeking test procedure during which cues were presented response-independently, and lever pressing produced no cues and no sucrose. Average (± SEM) active lever presses in **(B)** females and **(C)** males, and inactive lever presses in **(D)** females and **(E)** males. Average (± SEM) latency to initiate the first active lever press as a function of cue type and trial in **(F)** females and **(G)** males. # CS+ active lever pressing in Males > Females. ^CS+ > No cue, and CS−. ∇ First DS+ presentation < first presentation of CS+, DS−, No cue, and CS−. Schematic in **(A)** adapted from Servonnet et al. (2020).

### LY379268 effects on cue-induced sucrose seeking

Following the cue-induced sucrose seeking test, we determined the effects of the mGlu_2/3_ receptor agonist, LY379268 on cue-induced sucrose seeking. Rats received 3 cue-induced sucrose seeking tests (50 min/test; illustrated in Fig. 6A) as described above, except that LY379268 (0.3, 1.0 mg/kg) or saline (counterbalanced across tests) was administered s.c. 30 min before testing (Baptista & Weiss, 2004; Garceau et al., 2023).

**Figure 6:**
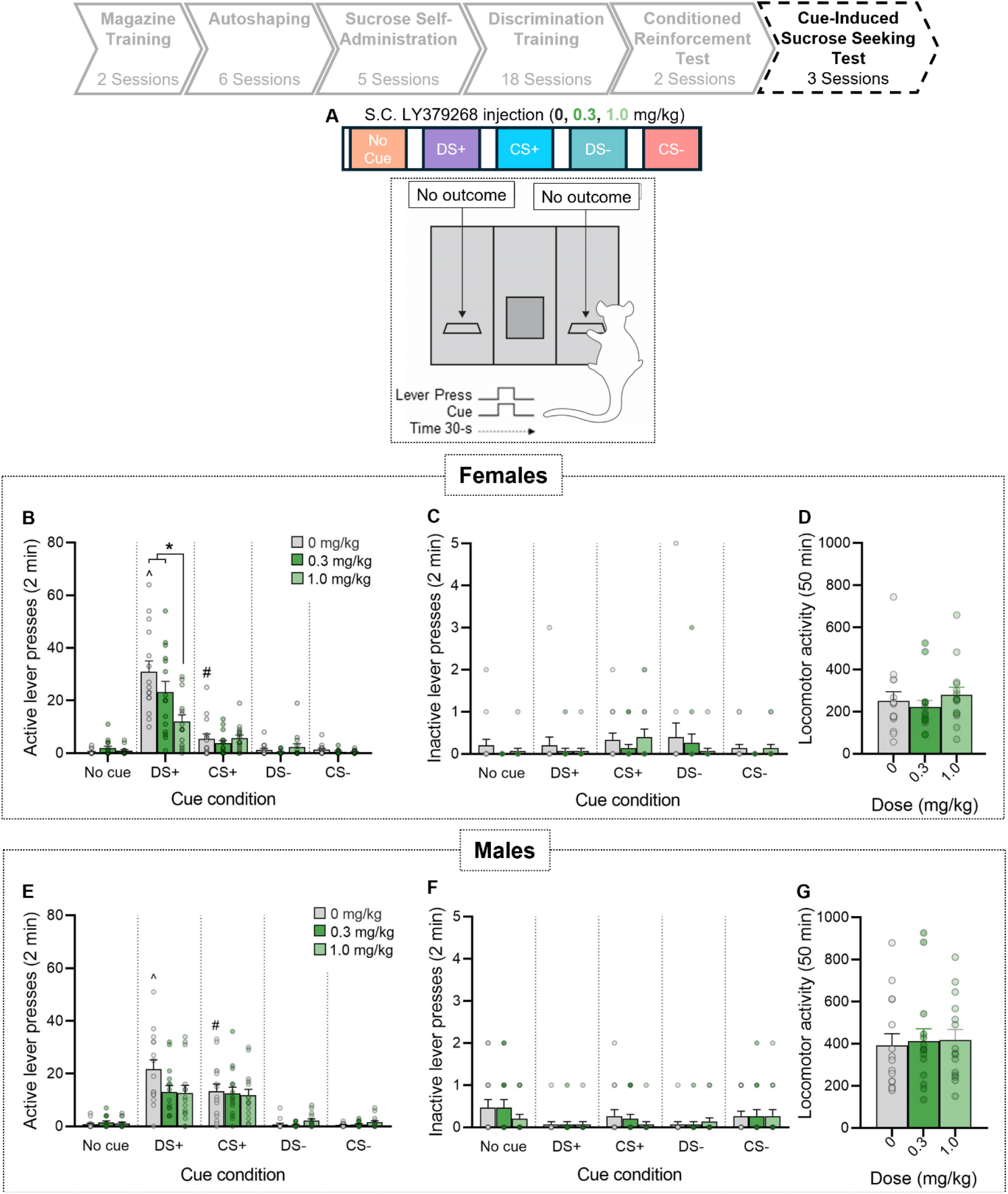
Effects of LY379268 on cue-induced sucrose seeking. **(A)** Schematic of the cue-induced sucrose seeking test procedure during which cues were presented response-independently, and lever pressing produced no cues and no sucrose. Average (± SEM) **(B)** active lever presses, (**C**) inactive lever presses and **(D)** locomotor activity in females. Average (± SEM) **(E)** active lever presses, (**F**) inactive lever presses and **(G)** locomotor activity in males. ^Under 0 mg/kg DS+ > CS+, DS− and No cue. # Under 0 mg/kg CS+ > CS−, and No cue. Schematic in **(A)** adapted from Servonnet et al. (2020).

### D-amphetamine effects on cue-induced sucrose seeking

Following the LY379268 tests, we determined the effects of D-amphetamine on cue-induced sucrose-seeking behaviour. To this end, rats received 2 cue-induced sucrose seeking tests (50 min/test; illustrated in Fig. 7A) as described above. Immediately before each test, rats received saline or 0.75 mg/kg D-amphetamine (counterbalanced across tests) s.c. (Evenden & Robbins, 1985; Slezak et al., 2018).

**Figure 7:**
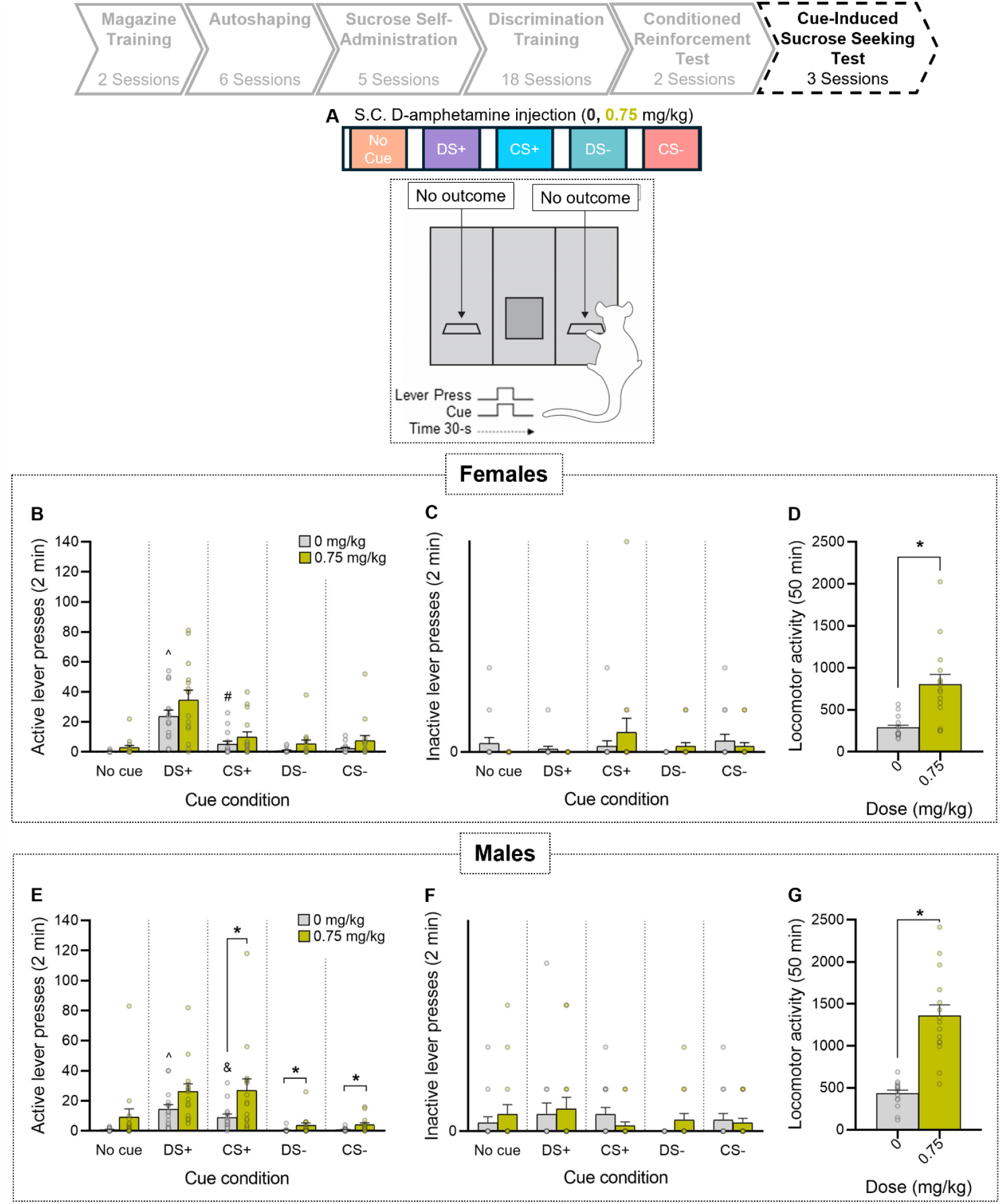
D-Amphetamine effects on cue-induced increases in sucrose-seeking behaviour. **(A)** Schematic of the cue-induced sucrose-seeking test procedure during which cues were presented response-independently, and lever pressing produced no cues and no sucrose. Average (± SEM) (**B**) active lever presses (**C**) inactive lever presses and (**D**) locomotor activity in females. Average (± SEM) (**E**) active lever presses (**F**) inactive lever presses and (**G**) locomotor activity in males. ^Under 0 mg/kg DS+ > CS+, DS− and No cue. # Under 0 mg/kg CS+ > No cue. & Under 0 mg/kg CS+ > No cue, and CS−. Schematic in **(A)** adapted from Servonnet et al. (2020).

## Statistical analyses

Data were analyzed using GraphPad Prism 10.2.3 and IBM SPSS 29.0.1, with an alpha level set at *p* < 0.05. We used two-sided p values except when a priori predictions were made, in which case we used one-sided p values. We used the Shapiro-Wilk test to assess data normality. When data were not normally distributed, we used nonparametric tests. We used Mann-Whitney tests to compare between-subjects factors (i.e., sex) and Wilcoxon signed-rank tests to compare within-subjects factors (i.e., active vs. inactive lever pressing, cue condition, treatment condition). When data were normally distributed, we conducted parametric tests (mixed ANOVA and *t*-tests) and the Greenhouse-Geisser correction to adjust for lack of sphericity where appropriate. We used Pearson’s correlation coefficient r to assess relationships between Pavlovian conditioned phenotype and *i)* days to meet acquisition criteria for discrimination training, *ii)* responding during a test for conditioned reinforcement, and *iii)* responding during a test for cue-induced sucrose seeking. We corrected for multiple comparisons using the Holm-Bonferroni method.

## Results

### Autoshaping

Figs. 1B and E illustrate average Pavlovian conditioned response bias across autoshaping sessions in sign-trackers (STs), goal-trackers (GTs), and intermediate (INTs). As illustrated by the pie chart insets in Figs. 1B and E, by the last session, 53% of females and 40% of males were classified as STs, 13% of females and 33% of males as INTs, and 33% of females and 27% of males as GTs. Across the sexes, GTs had high rates of magazine entries (Figs. 1C and F), and STs had high rates of lever contacts (Figs. 1D and G) in response to the lever CS. Interestingly, female INTs exhibited low levels of magazine entries, similar to female STs (Fig. 1C), and low levels of lever contacts (Fig. 1D), whereas male INTs showed high rates of both responses. In summary, ST was the most common Pavlovian conditioned phenotype in both sexes.

Table 1 shows that there were no statistically significant correlations between Pavlovian conditioned response bias (i.e., a sign- or goal-tracking phenotype) and active lever pressing during the conditioned reinforcement test (results described below), except for a positive correlation between response bias and responding for the DS+ in female rats (*r* = 0.64, *p* = 0.01). This indicates that during a test for conditioned reinforcement, females with a sign-tracking showed more vigorous instrumental pursuit of a sucrose-associated DS+. In parallel, there were no statistically significant correlations between Pavlovian conditioned response bias and active lever pressing behaviour during the cue-induced sucrose seeking test (results described below). Thus, a sign-vs. goal-tracking phenotype did not reliably predict DS+ or CS+ effects on sucrose-seeking behaviour.

**Table 1.**
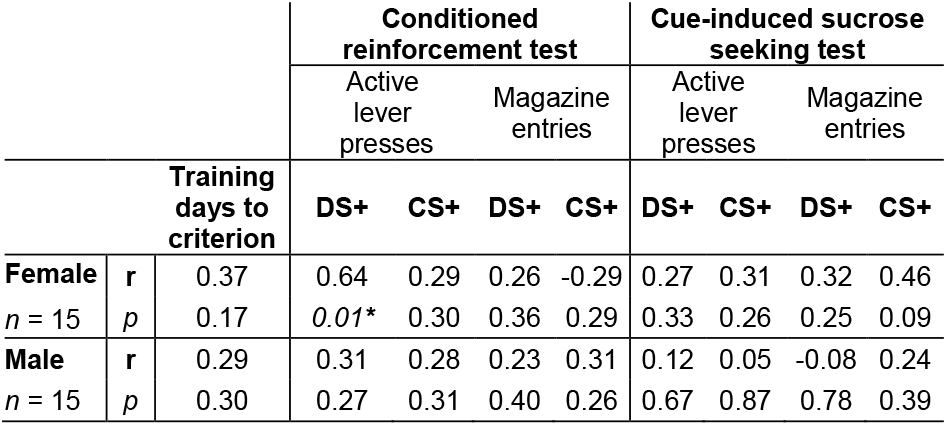
Correlations between Pavlovian conditioned response bias and *i)* days to reach discrimination training criteria, responding during *ii)* conditioned reinforcement test 1, and *iii)* cue-induced sucrose seeking test.

### Sucrose self-administration training

During FR1 and FR3 sessions (data not shown), females and males showed similar rates of active lever pressing, inactive lever presses, reinforcers earned, and magazine entries (see Table 2 for statistics). Collapsed across sex, rats pressed significantly more on the active vs. inactive lever (FR1, Session 1, *z* = −4.78, *p* = 0.003; Session 2, *z* = −4.79, *p* = 0.002; Session 3, *z* = −4.79, *p* = 0.002; FR3, Session 1, *z* = −4.78, *p* = 0.001; Session 2, *z* = −4.78, *p* = 0.001). Thus, both females and males learned to self-administer sucrose and retrieve it from the magazine.

**Table 2.**
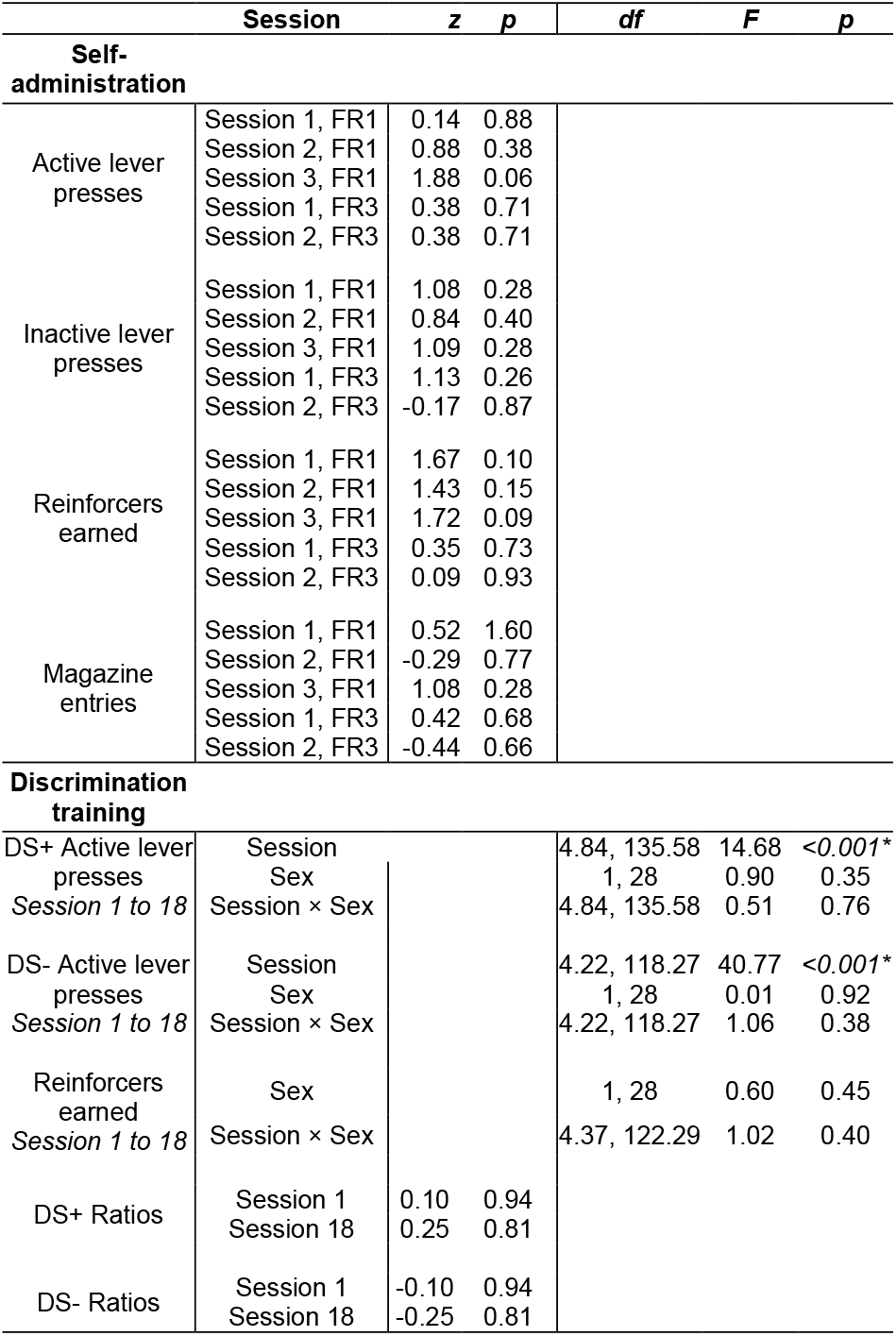
Lack of sex effects on behavioural measures recorded during sucrose self-administration and discrimination training.

### Sucrose self-administration under the control of discriminative stimuli

There were no sex differences in responding during discrimination training sessions (see Table 2 for statistics). Indeed, from Session 1 to 18 of discrimination training, females and males showed similar rates of active lever presses during DS+ and DS−presentation, number of sucrose reinforcers, and discrimination ratios during Session and Session 18. Therefore, data were pooled across the sexes in subsequent analyses.

Visual inspection of Fig. 2B shows that on Session 1, active lever presses peaked during DS−trials, during which rats made 70% of their active lever presses (Fig. 2B inset). By Session 15 (Fig. 2C), however, this pattern reversed, and active lever presses now peaked during DS+ trials, during which rats made 80% of their active lever presses (Fig. 2C inset). During sessions 16-18, in which the sequence of DS+ and DS−presentation was pseudo-randomized and the inter-trial interval length was varied, rats still suppressed responding during DS−trials and ramped up their responding during DS+ trials, making 80-81% of their active lever presses during the DS+ (Figs. 2D, E and insets). This indicates that rather than keeping time, the rats were relying on the discriminative stimuli to guide their sucrose self-administration behaviour.

To further highlight the control of self-administration behaviour by the DSs, Fig. 2F shows discrimination ratios for the DS+ and DS−. On Session 1, average discrimination ratio was significantly higher for the DS−vs. the DS+ (*z* = −4.72, *p* < 0.001), indicating that rats were pressing most on the active lever during DS−trials. However, this pattern of responding gradually reversed, such that on Session 18, where the discrimination ratio was greatest for the DS+ vs. the DS− (*z* = −4.67, *p* < 0.001). Moreover, from Session 1 to 18, the DS+ discrimination ratio significantly increased (*z* = 4.76, *p* < 0.001), while the DS−discrimination ratio significantly decreased (*z* = −4.76, *p* < 0.001). Accordingly, rats earned an increasing number of sucrose reinforcers across sessions (Fig. 2G; Session: *F* (4.37, 122.29) = 23.12, *p* < .001).

In summary, female and male rats learned to discriminate between a DS+ signaling sucrose availability and a DS−signaling reinforcer unavailability, such that they came to load up on sucrose during DS+ trials and inhibit responding during DS−trials.

### Tests for conditioned reinforcement

#### Test 1. All cue conditions tested during a single session

To assess the conditioned reinforcing properties of the DS+ and CS+, all rats received a test during which responding on the active lever produced the DS+, CS+, DS−, CS−, or No Cue for 5 s. Females (Fig. 3B) and males (Fig. 3C) showed similar rates of active lever presses during the No Cue (*z* = −0.02, *p* = 1.0), DS+ (*t*_(28)_ = −2.05, *p* = 0.20), DS−(*z* = 0.21, *p* = 0.87), and CS− (*z* = 0.63, *p* = 0.57) conditions. Of note, compared to females, males responded significantly more for the CS+ (*z* = −3.36, *p* < 0.001). Therefore, subsequent analyses are separated by sex.

Females made significantly more active than inactive lever presses (Figs. 3B, D) during the No Cue (*z* = −2.38, *p* = 0.03), DS+ (*z* = −2.94, *p* = 0.008), CS+ (*z* = −2.68, *p* = 0.02, and CS− (*z* = −2.68, *p* = 0.02) conditions. However, there was no significant lever discrimination for the DS−(*z* = −1.79, *p* = 0.07). Females also responded at higher rates on the active lever to earn DS+ presentation compared to all other cue conditions (Fig. 3B; vs. No Cue, *z* = −2.76, *p* = 0.01; CS+, *z* = −2.59, *p* = 0.02; DS−, *z* = −2.93, *p* = 0.005). In contrast, rats responded similarly on the active lever when this produced the CS+ vs. the CS− (*z* = 0.18, *p* = 0.43) or the No Cue condition (*z* = 0.10, *p* = 0.46). Thus, female rats did not respond more for a stimulus previously paired with sucrose delivery (CS+) than they did for a stimulus explicitly unpaired from sucrose delivery (CS−), suggesting that the CS+ did not acquire significant conditioned reinforcing value. In contrast, females showed high rates of responding for the DS+, suggesting that a discriminative stimulus previously signaling sucrose availability acquired strong conditioned reinforcing value.

Males made significantly more active compared to inactive lever presses (Figs. 3C, E) during the DS+ (*z* = −3.18, *p* = 0.004), CS+ (*z* = −3.18, *p* = 0.003), and No Cue (*z* = −2.53, *p* = 0.04) conditions. However, there was no significant lever discrimination for the DS−(*z* = −1.45, *p* = 0.15) or CS− (*z* = −0.99, *p* = 0.32). Similar to the females, males responded most on the active lever for the DS+ compared to the other cue conditions (Fig. 3C; vs. No Cue, *z* = 3.18, *p* = 0.003; DS−, *z* = −3.19, *p* = 0.001). In addition, and unlike the females, males made more active lever presses for the CS+ compared to the CS− (*z* = −3.30, p = 0.001) and No Cue (*z* = −3.30, p = 0.001) conditions, and had similar rates of responding for the CS+ and DS+ (*z* = −1.67, *p* = 0.10). Thus, in male rats, the DS+ and CS+ acquired equally strong conditioned reinforcing properties. In summary, when all cue conditions are made available during a single test session, the DS+ elicited the highest rate of instrumental responding across the sexes. In comparison, the CS+ was a strong conditioned reinforcer in male rats but was ineffective in females.

#### Test 2. One cue condition available per test session

To explore this apparent sex effect further, we gave rats additional conditioned reinforcement tests during which the CS+ and DS+ were presented on separate sessions (Fig. 4A).

Females made significantly more active vs. inactive lever presses (Fig. 4B, D) for the DS+ (*z* = −3.30, *p* = 0.001) and CS+ (*z* = −3.07, *p* = 0.002). There was no lever discrimination during the No Cue condition (*z* = −1.60, *p* = 0.09). Females again responded more vigorously on the active lever for the DS+ compared to either the CS+ (*z* = −2.54, *p* = 0.01) or No Cue (*z* = −3.24, *p* = 0.002) conditions. However, in contrast to what was observed in Test 1 (Fig. 3B), female rats pressed more on the active lever for the CS+ than they did for the No Cue condition (Fig. 4B; *z* = 2.13, *p* = 0.03). Thus, when female rats can lever press for a single cue type per session, they respond to obtain both a stimulus that previously signaled sucrose availability (DS+) and a stimulus previously paired with sucrose delivery (CS+). However, they show more vigorous instrumental pursuit of the DS+ vs. the CS+, suggesting stronger conditioned reinforcing value.

Males made significantly more active than inactive lever presses (Fig. 4C, E) during the DS+ (*z* = −3.11, *p* = 0.002), CS+ (*z* = −3.11, *p* = 0.003), and No Cue (*z* = −3.07, *p* = 0.002) conditions. Compared to the No Cue condition, male rats also showed greater rates of active lever pressing when this produced the DS+ (*z* = −3.11, *p* = 0.003) and CS+ (*z* = 2.97, *p* = 0.003). Males again showed similar rates of active lever pressing for the DS+ and the CS+ (*z* = −1.82, *p* = 0.07). Thus, male rats showed similar instrumental pursuit of a DS+ and a CS+.

In contrast to Test 1, here, females (Fig. 4B) and males (Fig. 4C) showed similar rates of responding for the CS+ (*z* = −1.50, *p* = 0.14). Females and males also responded similarly during the DS+ and No Cue conditions; respectively (*t*_(26)_ = −1.42, *p* = 0.17, and *z* = −1.07, *p* = 0.29). In summary, biological sex and test conditions interact to determine the conditioned reinforcing properties of sucrose-associated CSs. Specifically, when rats have access to both cue types during a test session, the CS+ is effective in reinforcing instrumental responding in males but not in females. In contrast, when rats have access to a single cue type during a test session, the CS+ and the DS+ significantly reinforce responding in both sexes. Importantly, in female rats, the DS+ gained more conditioned reinforcing value compared to the CS+, supporting higher rates of instrumental pursuit.

### Cue-induced sucrose seeking

We next assessed the extent to which response-independent presentations of the DS+ and CS+ triggered increases in sucrose-seeking behaviour, relative to baseline responding, during a test where lever pressing produced no cues or sucrose. As Figs. 5B, C show, there were no significant sex differences in rates of active lever pressing during No Cue (*z* = −1.45, *p* = 0.22), DS+ (*z* = −1.80, *p* = 0.07), DS−(*z* = −1.14, *p* = 0.29), or CS− (*z* = −1.52, *p* = 0.16) trials. However, males made significantly more active lever presses than did females during CS+ presentations (*z* = −3.74, *p* < 0.001), thus data were separated by sex in subsequent analyses.

Females pressed significantly more on the active versus inactive lever (Figs. 5B-D) only during DS+ (*z* = −3.30, *p* = 0.003) trials. There was no significant lever discrimination during the CS+ (*z* = −2.41, *p* = 0.07), DS−(*z* = −2.04, *p* = 0.08), CS− (*z* = −2.27, *p* = 0.07), or No Cue (*z* = −1.23, *p* = 0.22) trials. Importantly, cue type determined the vigor of responding on the active lever (Fig. 5B). The DS+ triggered higher rates of active lever pressing compared to the other cue conditions (No Cue, *z* = −3.30, *p* = 0.002; DS−, *z* = −3.30, *p* = 0.001) including the CS+ (*z* = −3.30, *p* = 0.001). As for the CS+, it evoked similar greater rates of active lever pressing compared to the CS− (*z* = −2.09, *p* = 0.04) and similar rates of responding compared to the No Cue condition (*z* = 1.90, *p* = 0.17). Thus, in female rats, only a stimulus previously signaling sucrose availability (DS+) significantly triggered increases in sucrose-seeking behaviour.

Males pressed significantly more on the active versus inactive lever (Figs. 5C-E) during DS+ (*z* = −3.41, *p* = 0.002), CS+ (*z* = −3.41, *p* = 0.003), and CS− (*z* = −2.40, *p* = 0.049) presentations. There was no lever discrimination during DS−(*z* = −1.97, *p* = 0.98), or No Cue (*z* = −1.73, *p* = 0.08) presentation. The DS+ triggered higher rates of responding on the active lever in male rats compared to the other cue conditions (Fig. 5C; vs. No Cue, *z* = −3.41, *p* = 0.001; DS−, *z* = −3.41, *p* = 0.001), but not compared to the CS+ (*z* = −1.85, *p* = 0.07). The CS+ evoked more active lever pressing than did the No Cue (*z* = −3.41, *p* = 0.002) and CS− (*z* = −3.42, *p* = 0.001) conditions. Thus, in male rats a stimulus signaling sucrose availability (DS+) and a stimulus predicting sucrose delivery (CS+) were similarly effective in increasing sucrose-seeking.

There were no sex differences in the total latency to make the first active lever press under the different cue conditions (data not shown; No Cue, *z* = 0.72, *p* = 0.78; DS+, *z* = 0.98, *p* = 1.00; CS+ *z* = 0.50, *p* = 0.26; DS−, *z* = 0.39, *p* = 0.44; and CS−, *z* = 0.84, *p* = 0.87). However, because active lever presses were different across the sexes, the data remained separated by sex for subsequent analyses.

Each cue condition was presented on 4 trials during the test and Figs. 5F-G show the latency to initiate active lever pressing as a function of trial number and cue condition in female and male rats, respectively. Female rats initiated lever pressing faster on the first vs. last presentation of each cue type (Fig. 5F; DS+, *z* = 2.82, *p* = 0.02; DS−, *z* = 2.52, *p* = 0.047; CS−, *z* = 2.52, *p* = 0.04), except under the No Cue (*z* = 2.02, *p* > 0.05) and CS+ (*z* = 2.09, *p* = 0.07) conditions. Cue type also determined the latency to the first lever press during the first cue presentation. Female rats initiated pressing fastest when presented with the DS+ compared to other cue conditions (Fig. 5F; vs. DS−*z* = 3.08, *p* = 0.006; CS+, *z* = 2.29, *p* = 0.04; No cue, *z* = 2.97, *p* = 0.012). In comparison, they had a similar latency to respond during the first presentation of the CS+ vs. the No Cue (*z* = 1.80, *p* = 0.07) or CS− (*z* = 1.77, *p* = 0.24) conditions. Thus, female rats initiated sucrose seeking faster when they encountered the DS+ but not the CS+.

Male rats initiated lever pressing faster on the first vs. last DS+ (Fig. 5G; *z* = 3.04, *p* = 0.01) and CS+ (*z* = 2.60, *p* = 0.04) presentation. In contrast, the delay to respond was similar on the first vs. last presentation of the No Cue (*z* = 2.20, *p* = 0.08), DS−(*z* = 1.86, *p* = 0.06), and CS− (*z* = 2.20, *p* > 0.05). As in the females, cue type determined the latency to initiate responding on the first cue presentation. Males initiated responding faster on the first presentation of the DS+ compared to the DS−(Fig. 5G; *z* = 3.04, *p* = 0.007) and No Cue (*z* = 3.06, *p* = 0.009) conditions. Also similar to females, males responded with a comparable latency to the first presentation of the CS+ vs. No Cue (*z* = 2.05, *p* = 0.06) and CS− (*z* = 1.69, *p* = 0.09) conditions. However, in contrast to females, males responded to the first DS+ and CS+ presentation with a similar latency (*z* = 1.57, *p* = 0.06). Thus, male rats initiated sucrose-seeking behaviour at a similar speed when they encountered the DS+ and CS+.

### LY379268 effects on cue-induced sucrose seeking

During a test where sucrose was unavailable, we evaluated the effects of the mGlu_2/3_ receptor agonist LY379268 (0, 0.3, and 1 mg/kg) on the ability of response-independent cue presentations to prompt increases in sucrose-seeking behaviour.

Under saline conditions, female rats pressed significantly more on the active vs. inactive lever (Fig. 6B, C) in response to DS+ (*z* = −3.41, *p* = 0.002), CS+ (*z* = −2.59, p = 0.04), and CS− (*z* = −2.51, *p* = 0.04) presentations, while there was no significant lever discrimination during No Cue (*z* = −0.92, *p* = 0.36) and DS−(*z* = −1.36, *p* = 0.18) conditions.

Female rats also responded at higher rates on the active lever during DS+ presentations compared to the other cue conditions (Fig. 6B; No Cue, *z* = −3.41, *p* = 0.001; DS−, *z* = −3.41, *p* = 0.001; CS+, *z* = −3.41, *p* = 0.001). Similarly, CS+ presentations evoked more responding than the No Cue (*z* = −2.65, *p* = 0.024) and CS− (*z* = −2.17, *p* = 0.03) conditions. LY379268 dose-dependently reduced the increases in active lever pressing triggered by the DS+ (Fig. 6B). At 1 mg/kg, LY379268 significantly decreased responding to the DS+ compared to both saline (*t*_(14)_ = −3.51, *p* = 0.01) and 0.3 mg/kg (*t*_(14)_ = −2.67, *p* = 0.04). Under 1 mg/kg, rats still made more active lever presses during the DS+ compared to the No Cue condition (*z* = 2.97, *p* = 0.006), suggesting that LY379268 decreased the ability of the DS+ to trigger peaks in sucrose-seeking behaviour without completely abolishing this effect. No other comparisons were statistically significant. LY379268 did not significantly influence locomotion in females (Fig. 6D; 0.3 mg/kg vs. saline, *z* = −0.54, *p* = 0.29; 1 mg/kg vs. saline, *z* = 0.82, *p* = 0.41; 0.3 mg/kg vs. 1 mg/kg, *z* = −1.85, *p* = 0.10). Thus, in female rats, activating mGlu_2/3_ receptors reduced the increased sucrose seeking prompted by the DS+, without producing significant locomotor effects.

Under saline conditions, male rats pressed significantly more on the active versus inactive lever (Fig. 6E, F) in response to DS+ (*z* = −3.30, *p* = 0.002) and CS+ (*z* = −3.18, *p* = 0.006) presentations. There was no significant lever discrimination under No Cue (*z* = −0.42, *p* = 0.67), CS− (*z* = −1.41, *p* = 0.16), or DS− (*z* = −1.07, *p* = 0.29) conditions. Males also responded at higher rates on the active lever (Fig. 6E) during DS+ presentations compared to the No Cue (*z* = −3.30, *p* = 0.002), DS− (*z* = −3.30, *p* = 0.001), and CS+ (*t*_(14)_ = −2.52, *p* = 0.03) conditions. The CS+ also evoked more active lever pressing than did the No Cue (*z* = −3.30, *p* = 0.001) and CS− (*z* = −3.18, *p* = 0.001) conditions. In contrast to females, LY379268 did not significantly influence the increase in sucrose-seeking behaviour triggered by the DS+ (Fig. 6E) (0.3 mg/kg vs. saline, *z* = −1.65, *p* > 0.05; 1.0 mg/kg vs. saline, *z* = −1.68, *p* = 0.14; 0.3 vs. 1.0 mg/kg, *z* = 0.40, *p* > .05). Similarly, LY379268 did not significantly influence responding to the CS+ (0.3 mg/kg dose vs. saline, *t*(14) = 0.16, *p* = 0.88; 1.0 mg/kg vs. saline, *t*_(14)_ = −0.41, *p* = 0.69; 0.3 mg/kg vs. 1.0 mg/kg, *t*_(14)_ = 0.40, *p* = 0.69). LY379268 also did not significantly influence locomotion in males (Fig. 6G; 0.3 mg/kg vs. saline, *z* = −0.23, *p* = 0.41; 1 mg/kg vs. saline, *z* = 0.77, *p* = 0.66; 1 mg/kg vs. 0.3 mg/kg vs. 1 mg/kg, *z* = −0.57, *p* = 0.57). Thus, mGlu_2/3_ receptor activation did not significantly influence cue-induced sucrose-seeking behaviour in male rats.

### D-amphetamine effects on cue-induced sucrose seeking

During a test where sucrose was unavailable, we evaluated the effects of D-amphetamine (0 and 0.75 mg/kg) on the ability of response-independent cue presentations to prompt increases in sucrose-seeking behaviour.

Under saline conditions, females pressed significantly more on the active vs. inactive lever (Fig. 7B vs. C) in response to DS+ (*z* = −4.71, *p* < 0.001), CS+ (*z* = −3.69, *p* = 0.001), No Cue (*z* = −2.41, *p* = 02), DS− (*z* = −3.33, *p* = 0.002), and CS− (*z* = −3.74, *p* = 0.001) conditions. Females showed higher rates of active lever pressing when presented with the DS+ vs. the other cue conditions (Fig. 7B; vs. No Cue, *z* = −3.41, *p* = 0.001; DS−, *z* = −3.41, *p* = 0.001; CS+, *z* = −3.18, *p* = 0.001). The CS+ also evoked active lever pressing compared to the No Cue condition (*z* = −2.61, *p* = 0.03), but not the CS− (*z* = −0.91, *p* = 0.36). Regardless of cue condition, D-amphetamine did not significantly influence active lever pressing compared to saline (No Cue, *z* = 2.04, *p* > 0.05; CS−, *z* = 1.34, *p* = 0.18; DS−, *z* = 2.37, *p* = 0.09; DS+, *z* = 1.40, *p* = 0.08; CS+, *z* = 1.19, *p* = 0.12). However, D-amphetamine significantly enhanced total locomotor activity (Fig. 7D; z = 3.41, *p* < 0.001). Thus, in female rats, D-amphetamine did not influence the increase in sucrose-seeking behaviour triggered by the DS+.

Under saline conditions, male rats pressed significantly more on the active vs. inactive lever (Fig. 7E vs. F) in response to DS+ (*z* = −4.78, *p* < 0.001),CS+ (*z* = −4.71, *p* < 0.001), No Cue (*z* = −2.97, *p* = 0.006), DS− (*z* = −2.95, *p* = 0.003), and CS− (z = −3.40, *p* = 0.002) conditions. Male rats also showed the highest rates of active lever pressing during DS+ presentations (Fig. 7E; vs. No Cue, *z* = −3.41, *p* = 0.001; DS−, *z* = −3.41, *p* = 0.002; CS+, *z* = −2.49, *p* = 0.013). The CS+ also evoked more active lever pressing compared to the No Cue (*z* = −3.18, *p* = 0.002) and CS− (*z* = −3.30, *p* = 0.002) conditions. D-amphetamine increased rates of active lever pressing (Fig. 7E) during the CS+ (*z* = 2.67, *p* = 0.02; DS−, *z* = 2.57, *p* = 0.03; CS−, *z* = 2.84, *p* = 0.02), however, the DS+ (*z* = 1.82, *p* > 0.05) and No Cue (*z* = 2.17, *p* = 0.06) conditions remained unaffected. As in the females, D-amphetamine significantly increased locomotion (Fig. 7G; *z* = 3.41, *p* < 0.001). Thus, in male rats, D-amphetamine increased sucrose-seeking responses indiscriminately across all cue types.

## Discussion

We developed a new trial-based procedure to compare reward-seeking behaviour triggered by conditioned stimuli (CSs) and discriminative stimuli (DSs) in the same subjects during the same session. Female and male rats learned to self-administer sucrose paired with a CS+ during discrete DS+ trials that signaled sucrose availability, and to inhibit responding during DS−trials that signaled sucrose unavailability. The rats maintained this pattern of responding even when both the order of DS+ and DS−presentation and the duration of inter-trial intervals became unpredictable during each session. Thus, our new procedure enables DSs to gain excitatory and inhibitory properties, such that responding comes under the control of the DSs (see also Ndiaye et al., 2024). Subsequently, when tested in the same session and in the absence of sucrose, the DS+ and CS+ equally reinforced instrumental responding in male rats. However, only the DS+ acted as a conditioned reinforcer in females. When tested in separate sessions, both the CS+ and DS+ reinforced responding in males, while in females, the CS+ now became a conditioned reinforcer, though the DS+ remained more effective. Also in the absence of sucrose, response-independent presentations of the DS+ prompted increases in sucrose seeking in both sexes, while the CS+ was effective only in males. Systemic injections of LY379268, a selective mGlu_2/3_ receptor agonist, suppressed the DS+-induced increases in sucrose seeking in females, with no effect in males. Finally, systemic D-amphetamine non-specifically enhanced sucrose seeking across cue types in males, with no effect in females.

### A new discrimination procedure

Our procedure builds on decades of research investigating the influence of DSs on reward-seeking behaviour. Typical approaches involve presenting the DSs signaling reward availability (DS+) and unavailability (DS−) in separate training sessions and/or over prolonged periods of exposure (Ciccocioppo et al., 2001; Colwill & Rescorla, 1990; Hauser et al., 2023; Laque et al., 2019; Suto et al., 2016; Weiss et al., 2001; Zhao et al., 2006). While these methods provide insight into conditioned appetitive behaviour, they make it difficult to parse out the contributions of DSs from those of context. Moreover, DSs also occur as discrete events in real-world scenarios, making it important to also model the effects of discrete, trial-based DS presentations on reward pursuit. While trial-based discrimination tasks have been refined recently (Madangopal et al., 2019, 2021), they have focused on DSs exclusively, omitting CSs. Animals evolve in cue-rich environments where survival and reproduction depend on the ability to process several reward-associated cues at the same time. Our procedure allows rats to simultaneously form relationships between discriminative stimuli, conditioned stimuli, and reward, in single sessions. This design enables one to later determine how distinct cue types with unique associative meanings influence seeking behaviour within a single test session. Advantages of this test design include a reduced likelihood of extinction of stimulus-outcome and stimulus-response relationships, the assessment of individual differences in responding to conditioned versus discriminative cues, and increased compatibility with time-locked neural manipulation techniques, such as optogenetics and in vivo imaging. As such, our new behavioural procedure opens new avenues for exploring the mechanisms underlying the effects of different types of reward-associated cues on appetitive behaviour.

### Conditioned reinforcing properties of the DS+ and CS+

After discrimination training, rats could press a lever, previously associated with sucrose delivery, to now earn presentations of a DS+, CS+, DS−, CS−, or no cue. Regardless of whether the DS+ was tested alone or amidst other cue types, it supported high rates of instrumental responding across the sexes and (see also Collins et al., 2023; Ndiaye et al., 2024). In contrast, the conditioned reinforcing properties of the CS+ varied as a function of both biological sex and test condition. When rats had access to all cues in alternating trials within one session, the CS+ was an effective conditioned reinforcer of instrumental behaviour in males but not females. However, if rats could only produce one cue during the test session, the CS+ reinforced responding in both sexes, and males responded equally for the CS+ and DS+ while females responded most for the DS+. These findings are consistent with others showing that, in male rats, CSs acquire robust conditioned reinforcing properties, maintaining reward-seeking actions in the absence of the primary reward (Davis & Smith, 1976; de Wit & Stewart, 1981; Milton & Everitt, 2010; Nie & Janak, 2003; Robbins, 1978; Robinson & Berridge, 1993; Servonnet et al., 2020, 2023; Shaham et al., 2003; Shalev et al., 2002; Stewart, 1984). Notably, the DS+ had the strongest conditioned reinforcing effects in both sexes, and the CS+ had weaker effects in females.

Why might CS+ effects be weaker in females vs. males? This cannot be explained by differences in the degree of exposure to the sucrose reward or CS+/sucrose pairings, as the sexes did not differ on these measures during prior training. Using a different primary reinforcer (i.e., cocaine), some studies report no sex differences in rates of instrumental responding reinforced by a CS+ (Feltenstein et al., 2011; Kawa & Robinson, 2019; Lynch et al., 2005; Ndiaye et al., 2024; Pitchers et al., 2015). Yet others find that compared to male rats, female rats respond less to earn a drug-paired CS+ (Fuchs, Evans, Mehta, et al., 2005; Shalev et al., 2002). Females may attribute less motivational significance to reward-associated CSs via both estrous cycle-dependent and -independent differences in neurobiology (Fuchs, Evans, Mehta, et al., 2005). For instance, intrinsic sex differences in midbrain dopamine systems can contribute to sex differences in learning and motivational processes (Gillies et al., 2014; O’Connell & Hofmann, 2011). Whatever the underlying mechanisms, our findings underscore the importance of studying both female and male subjects to account for the natural diversity of brain function and dysfunction (Shansky, 2024).

### The ability of response-independent DS+ and CS+ presentation to trigger increases in sucrose-seeking behaviour

In addition to their conditioned reinforcing effects, reward-associated cues can act as response activators when presented response independently, both triggering and invigorating reward-seeking actions (Colagiuri & Lovibond, 2015; Dickinson & Dawson, 1987; Rescorla & Solomon, 1967). We found a sex effect in the ability of response-independent DS+ and CS+ presentation to boost sucrose-seeking responses in the absence of sucrose reward. In females, the DS+ was consistently more effective than the CS+. In males, however, the DS+ and CS+ were initially equally effective in enhancing sucrose-seeking responses, but with repeated testing, the DS+ emerged as being more effective than the CS+.

The ability of the DS+ to elicit peaks in sucrose seeking depended on its positive relationship with sucrose reward, because presentation of cues signaling the absence of reward (DS− and CS−) did not increase sucrose-seeking behaviour. In addition, the DS+ was presented independently of behaviour during cue-induced sucrose seeking tests. As such, the increased sucrose seeking produced by the DS+ does not involve the cue’s conditioned reinforcing properties, but likely its Pavlovian incentive motivational effects (Robinson & Berridge, 1993). These findings are consistent with others showing that when presented independently of the subject’s actions, a DS+ is superior in boosting reward seeking compared to a CS+ (Ciccocioppo et al., 2001; Collins et al., 2023; Deroche-Gamonet et al., 2002; Di Ciano & Everitt, 2003; Madangopal et al., 2019, 2021; Ndiaye et al., 2024; Pitchers et al., 2015; Weissenborn et al., 1995; Yun & Fields, 2003). Together, the findings suggest that through associative learning, cues indicating reward availability (DS+) may develop stronger incentive motivational properties than cues signaling reward delivery (CS+). This difference may enable DSs to elicit stronger cravings for the associated reward.

### Sign-trackers versus goal-trackers

Pavlovian conditioned phenotype generally did not predict responding evoked by, or reinforced by, the CS+ or DS+. This is inconsistent with previous studies suggesting that sign-trackers are more likely to respond to CSs, while goal-trackers respond more to DSs (Kuhn et al., 2022; Pitchers et al., 2017; Saunders et al., 2013; Saunders & Robinson, 2010; Yager & Robinson, 2010). Our findings are, however, in line with other studies reporting no strong predictive relationship between Pavlovian conditioned phenotype and CS vs. DS effects on behaviour (Chang et al., 2022; Kawa et al., 2016; Martin et al., 2022). Differences in methodology may explain the disparities between studies. For instance, in contrast to some studies where sucrose seeking-behaviour was extinguished before testing DS and CS effects (Pitchers et al., 2017; Saunders et al., 2013; Saunders & Robinson, 2010; Yager & Robinson, 2010), our rats did not receive any extinction. In previous work, the behavioural effects of CSs (Saunders & Robinson, 2010) and DSs (Pitchers et al., 2017) were also assessed separately, whereas in the present study this was done in the same subjects, within the same test session.

### Effects of mGlu_2/3_ receptor activation on DS+-triggered increases in sucrose seeking

Injection of a selective mGlu_2/3_ receptor agonist (LY379268) supressed the increase in sucrose seeking evoked by DS+ presentations in female rats. This extends the findings of Garceau et al (2023) where the same LY379268 doses that were used here also reduced increases in reward-seeking behaviour evoked by a CS+, using a Pavlovian-to-Instrumental transfer design. MGlu_2/3_ receptors are mainly extrasynaptic and located on presynaptic terminals where their activation reduces glutamate release into the synaptic cleft (Conn & Pin, 1997; Imre, 2007; Schoepp, 2001). Thus, our findings are consistent with prior work showing that both synaptic glutamate neurotransmission (Derman et al., 2020; Derman & Ferrario, 2018; Feltenstein & See, 2007; Garceau et al., 2021; Khoo et al., 2019; Lu et al., 2007; Malvaez et al., 2015) and activity at mGlu_2/3_ receptors (Bäckström & Hyytiä, 2005; Baptista & Weiss, 2004; Bossert, Gray, et al., 2006; Bossert, Poles, et al., 2006; Lu et al., 2007; Uejima et al., 2007) contribute to the ability of reward-associated cues to increase reward-seeking responses. Importantly, the LY379268 doses we tested did not change locomotor behaviour (but see Garceau et al., 2023). Thus, the ability of the agonist to suppress the increases in sucrose seeking prompted by the DS+ is not due to a general motor deficit. We propose that activation of mGlu_2/3_ receptors suppressed the conditioned incentive salience of the DS+, diminishing its ability to trigger the motivational state that enhances reward seeking (Wyvell & Berridge, 2000).

In apparent contrast to some other studies (Bäckström & Hyytiä, 2005; Baptista & Weiss, 2004; Bossert, Gray, et al., 2006; Bossert, Poles, et al., 2006; Garceau et al., 2023; Lu et al., 2007; Uejima et al., 2007), LY379268 had no significant effect on the increases in sucrose-seeking behaviour triggered by the DS+ or CS+ in male rats. However, some of these studies used LY379268 doses higher than those used here, and these doses also impaired spontaneous locomotor behaviour (Bäckström & Hyytiä, 2005; Baptista & Weiss, 2004). Other studies used cue-induced reward-seeking tasks different from those we used here (Bossert, Poles, et al., 2006; Garceau et al., 2023; Lu et al., 2007; Uejima et al., 2007), potentially engaging different psychological and neurobiological processes as well. It is not clear why LY379268 was effective in females only. This is unlikely to involve intrinsic sex differences in the drug’s pharmacological profile, because doses of LY379268 similar to ours can effectively suppress behavioural responses to reward cues in males (Bäckström & Hyytiä, 2005; Baptista & Weiss, 2004; Bossert, Gray, et al., 2006; Bossert, Poles, et al., 2006; Garceau et al., 2023; Lu et al., 2007; Uejima et al., 2007). This suggests that larger doses would not be necessary for males compared to females. Still, our findings can be explored in future studies comparing the effects of other mGlu_2/3_ receptor agonists on cue-triggered reward seeking across the sexes.

### D-amphetamine enhances sucrose-seeking behaviour in male rats only

Injection of the indirect dopamine agonist, D-amphetamine increased sucrose seeking in males across cue types (i.e., CS+, DS−, and CS−). D-amphetamine, administered systemically, enhances the reinforcing properties of CSs (Killcross et al., 1997) and D-amphetamine injected specifically into the nucleus accumbens shell also boosts sucrose seeking evoked by a CS (Wyvell & Berridge, 2000). These effects of D-amphetamine are thought to involve activation of mesolimbic dopamine neurotransmission, leading to enhancement of the incentive salience attributed to reward cues (Wyvell & Berridge, 2000). However, in our experiment, D-amphetamine effects in males were non-specific, increasing lever pressing for sucrose even when we presented cues that had consistently signaled the absence of sucrose reward (DS− and CS−). D-amphetamine also increased locomotor activity. Thus, one possibility is that the increase in lever pressing behaviour we observed resulted from the psychomotor activating effects of D-amphetamine. We used 0.75 mg/kg specifically because lower doses do not significantly influence the response to reward cues in otherwise naive rats (Bédard et al., 2011, 2013; Mead et al., 2004; Robbins, 1978; Robbins et al., 1983). Future studies can address this by testing smaller D-amphetamine doses.

There was no significant effect of D-amphetamine on cue-triggered increases in sucrose seeking in female rats. However, D-amphetamine also enhanced locomotion in females. We speculate that D-amphetamine’s psychomotor activating effects caused greater interference with lever-pressing behaviour in the females. Female rats can show a greater psychomotor response to lower doses of D-amphetamine compared to males (Becker et al., 1982). However, the caveats are that we only tested a single D-amphetamine dose and did not measure the full extent of the drug’s psychomotor activating effects (i.e., stereotypy). This can be addressed in future work.

## Conclusions

We developed a new trial-based procedure to assess the unique and similar effects of CSs and DSs on reward seeking in the same subjects within a single test session. Using this procedure, we find that – across the sexes – a DS+ signalling reward availability and a CS+ predictive of reward delivery act as a conditioned reinforcer of instrumental responding; however, the DS+ is significantly more effective than the CS+ in females. The DS+ is also more effective overall than a CS+ in triggering increases in reward-seeking actions, and an mGlu_2/3_ receptor agonist can suppress this DS+ effect in females only. We conclude that this new discrimination training and testing procedure can be used to examine the effects of different classes of reward-associated cues on brain and behaviour, while also allowing for the characterization of potential sex effects in these processes.

## Acknowledgements

We are grateful to Drs. Rajtarun Madangopal and Yavin Shaham for advice during the development of the behavioural procedures used in the present work. This research was funded by grants to A.-N.S. from the National Science and Engineering Research Council of Canada (Grant 355923) and from the Courtois Fund. S. N. is supported by a Master’s Research Scholarship from the Fonds de recherche du Québec - Nature and technologies Sector. M. R. L. is supported by a postdoctoral fellowship from the Natural Sciences and Engineering Research Council of Canada.

